# Recollection is more than retrieving context or source memory: evidence from ERPs of Recognition and Source Memory combinations

**DOI:** 10.1101/2020.10.14.339697

**Authors:** Richard J. Addante, Alana Muller, Lindsey A. Sirianni

## Abstract

The goal of this study was to investigate a relatively unstudied memory condition for paradoxical combinations of item + source memory confidence responses, which challenged the conventional views of the memory processes supporting item and source memory judgments. We studied instances in which people provided accurate source memory judgments (conventionally ascribed as representing recollection) after having first produced low-confidence item recognition hits for the same items (conventionally thought to reflect familiarity-based processing). This paradoxical combination does not fit traditional accounts of being recollection (because it had low-confidence recognition) nor accounts of familiarity (since it had accurate source memory), and event-related potentials (ERPs) were used to adjudicate which processes support these kinds of memories. ERP results were unlike the conventional ERP effects of memory, lacking both an FN400 and the parietal old-new effect (LPC), and instead exhibited a significant negative-going ERP effect occurring later in time (800-1200ms) in central-parietal sites. Behavioral measures of response times revealed a crossover interaction: low confident recognition hits were slower during recognition but faster during source memory when compared to the opposite pattern seen for instances of high confident hits. Results provide a comprehensive characterization of the individual variability of the FN400 and LPC effects of memory, while adding the behavioral and physiological characterization of a late negative-going ERP effect for accurate source memory without recollection. Conclusions indicated that episodic context could be retrieved independently from recollection, while suggesting a role for a process of context familiarity that is independent from item-familiarity.

**Highlights:**

- Recollection is often defined as remembering the source or context of information
- Prior work used ERPs to identify times when source memory did not have recollection
- Current work replicated ERPs with added response times and measures of variance
- Recollection was not evident in certain source memories, which had a negative ERP
- Recollection is independent of context and is more than just remembering sources

**Graphical Abstract:** 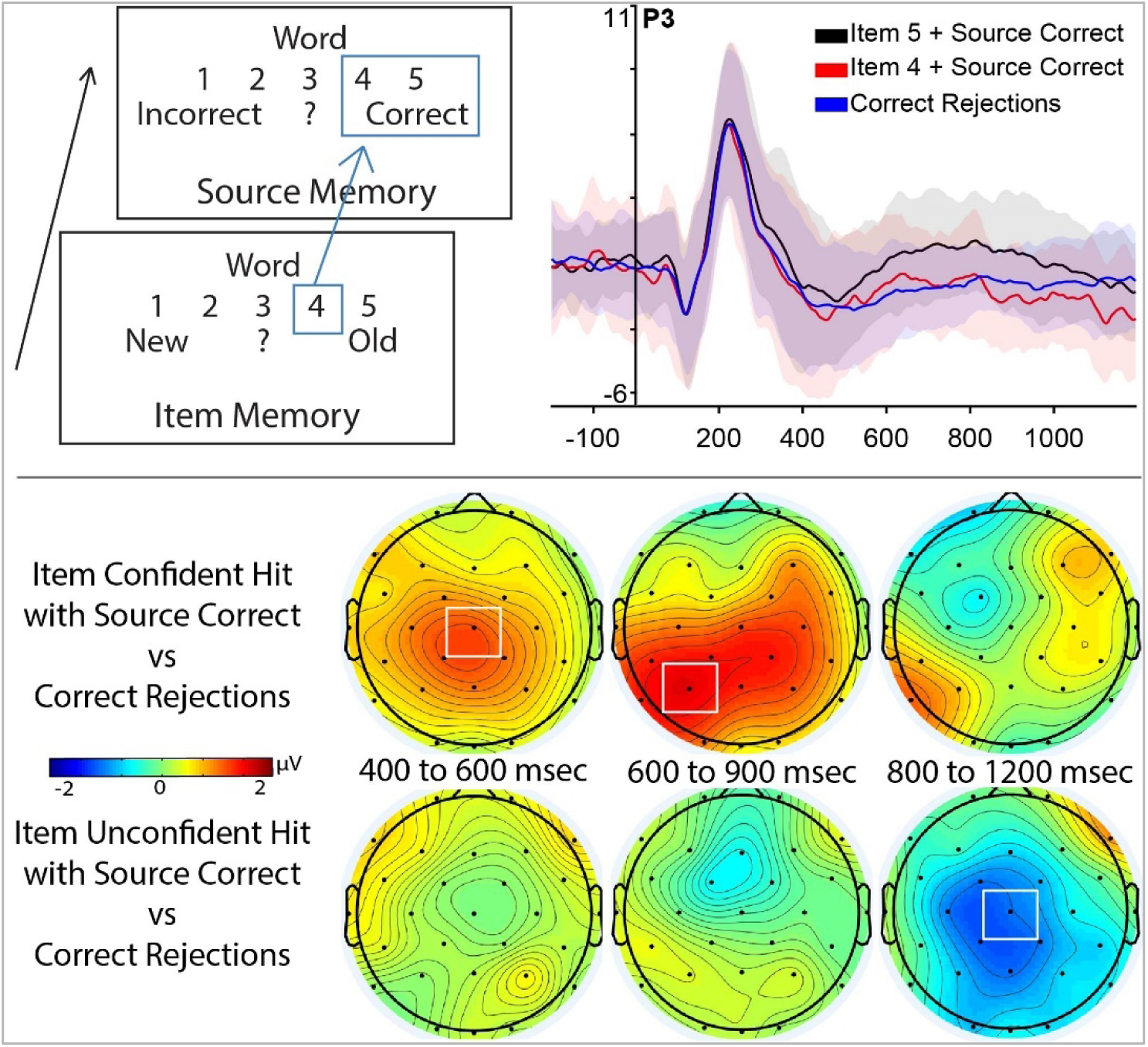

## 1.0 Introduction

In human long term memory, theoretical models of conscious episodic recognition are largely governed by the cognitive processes of familiarity and recollection for several decades (Diana, Yonelinas, & Ranganath, 2007; Eichenbaum, Yonelinas, & Ranganath, 2007; Mandler, 1980, 2008; Ranganath, 2010a, 2010b; Michael D. Rugg & Yonelinas, 2003; Squire & Wixted, 2011; Squire, Wixted, & Clark, 2007; Wixted, 2007; Wixted & Mickes, 2010; Yonelinas, 1994; Yonelinas, 2001; Yonelinas, 2002; Yonelinas, Aly, Wang, & Koen, 2010; Yonelinas, Ranganath, Ekstrom, & Wiltgen, 2019a)(Aktinson & Juola, 1974). Recollection can be operationalized as the declarative retrieval of episodic information of both the item and context bound together into a cohesive retrieval of the episodic event (for review see Diana et al., 2008), and is usually associated with the retrieval of contextual information surrounding the item of the event (Addante et al. 2012a; for reviews see Eichenbaum et al., 2007; Yonelinas et al. 2010; Ranganath 2010). The item in the event, however, may be retrieved without recollection and via reliance upon familiarity, typically conceptualized as retrieval of an item from a prior episode but without the associated contextual information in which it occurred. Familiarity occurs, for instance, when a person can remember that someone seems familiar from the past but cannot retrieve details of context, such as who the person is or from where they know them. Recollection, on the other hand, would be remembering precisely who someone is and the context of how you know them from a prior episode of one’s past experience.

Each of these two memory phenomena have been found to be dissociable cognitive processes (Yonelinas, 2002), with dissociable neural substrates in the medial temporal lobes (Ranganath et al., 2004), neuropsychologically dissociable among patient impairments (Addante, Ranganath, Olichney, & Yonelinas, 2012; Düzel et al., 1999; Mecklinger, von Cramon, & Matthes-von Cramon, 1998), and with distinct patterns of electrophysiology at the scalp that is both spatially and temporally dissociable in event-related potentials (ERPs) (Addante et al., 2012; Curran, 2000; Friedman, 2013; Gherman & Philiastides, 2015; Rugg et al., 1998; Rugg & Curran, 2007). Electrophysiologically, familiarity has been associated with ERP differences in memory trials during a negative-going peak at the mid-frontal scalp sites at approximately 400 milliseconds to 600 milliseconds post stimulus, called the mid-frontal old-new effect, or FN400 (for frontal-N400 effect). On the other hand, recollection has been associated with differences between memory conditions occurring at a peak in the ERP at the parietal region of the scalp from approximately 600 milliseconds to 900 milliseconds, often referred to as a late parietal component (LPC) (Addante et al., 2012; Leynes et al., 2005; for reviews see Rugg & Curran, 2007; Friedman, 2013).

Among studies of episodic recollection, ‘source memory’ has been one of the most reliable methods used to assess instances of recollection for episodic events in both behavioral and cognitive neuroscience measures, because it has traditionally been assumed that if someone remembers the source of information they obviously have retrieved information about the context of that memory and hence ‘must’ be reflecting recollection of that episode. As such, source memory has also been one of the most useful ways used to understand memory, amnesia deficits, and more. However, source memory measures remain often utilized in a relatively crude manner of simply comparing a basic conditions of source correct to incorrect source decisions, in order to represent times when people use recollection (Wilding & Rugg, 1996) (though for exceptions see (Ingram, Mickes, & Wixted, 2012), and it is both possible and likely that source memories are more complex than that simplistic dichotomy. For instance, other ways that researchers have investigated recollection and familiarity has been to use confidence scales for people to rate the extent of how well they remember something (Yonelinas, Otten, Shaw, & Rugg, 2005), and while familiarity has been considered to range across lower levels of confidence, recollection has generally been considered to be captured by responses of high confidence or correct source judgments.

A core goal of the current study was to investigate a relatively unstudied memory condition for paradoxical combinations of item + source memory responses, which had been reported earlier in a prior study (R. J. Addante, Ranganath, & Yonelinas, 2012) and challenged the conventional views of the roles for item and source memory processes. This unique condition had referred to instances in which people provided accurate source memory judgments (conventionally ascribed by researchers as representing recollection) despite having also first produced low-confidence item recognition hits (‘4’s’) for the same items (conventionally thought to reflect familiarity-based processing). This paradoxical combination could not be deciphered by traditional accounts of it being recollection (because despite the accurate source memory it had low-confidence recognition) nor by accounts of familiarity (since despite its low-confidence recognition it had accurate source memory), and ERPs were used to adjudicate which processes support these kinds of memories. The ERP results for this condition were unlike the conventional ERP effects of memory (they entirely lacked both an FN400 and LPC), and instead they exhibited a significant negative-going ERP effect occurring in later time (800-1200ms) in different places on the scalp (fronto-central sites). Since the findings were replicated in a second independent laboratory from that lab (R. J. Addante, Watrous, Yonelinas, Ekstrom, & Ranganath, 2011), authors concluded that context could be retrieved independently from recollection, which challenged theoretical models of memory that typically ascribed source memory to the act of recollecting.

The results reported in 2012 were novel, and not previously seen for their conditions, topography, or timing of ERP effects. Nevertheless, there remained several inherent limitations from its experimental design. For example, while the findings included observing the results replicate in a second experiment’s re-analysis of a prior study, these were studies from the same internal lab at UC Davis (USA), and with relatively small sample sizes (between N=12 and N=25) of a homogenous population and collected with the same researchers, recording devices, and protocols (BioSemi Systems). These findings were also limited for interpretations by common data reporting practices during that era of not reporting measures of variance for either the physiological signals or the behavioral measures of memory accuracy, which have taken prominence in the field in the time since then due to their omission’s potential to obscure the true nature of patterns in data (Rousselet, Foxe, & Bolam, 2016; Rousselet & Pernet, 2011). Given the novel findings thereof, we have since considered inclusion of such measures of variability as paramount for being able to assess the full scope and validity of the data results.

Furthermore, the prior findings maintained the limitation of not affording behavioral measures of reaction times for the critical conditions of memory that were compared; indeed, it lacked any reaction time measures at all. The lack of reaction time measures in the 2012 studies was due to the experimental design having been intentionally optimized to facilitate ERP data analysis that would be free from artifacts such as motion from responses, and thus instructed participants to withhold their motor responses of memory until after a 1500 msec window of time had passed for the mnemonic stimuli processing. In contrast, since the current study sought to specifically assess reaction time measures to help inform the cognitive theories of the processes underlying the physiological effects for the mnemonic conditions. As such, the study was designed to permit subjects to respond at their own pace of memory retrieval, and focused exclusion of any potential artifacts based on using artifact correction and rejection techniques such as independent component analysis (see Methods for further details).

The current investigation thus sought to extend that original finding of Addante et al. (2012) with a full-scale replication of the same paradigm assessing the extent to which it would replicate in a wholly independent laboratory (San Bernardino, California), using different recording hardware (ActiCHamp, as opposed to Biosemi used in the 2012 studies), different researchers collecting the data, and different demographic populations (77% non-Caucasian, 56% Latino in San Bernardino, vs % others in Davis, CA, respectively), while taking efforts to double the same size from the original N=25 reported in 2012 to the current sample of N=56 participants in the current study. Moreover, the current investigation sought to make substantive additions to prior findings by including measures of variance for data reporting that have been called for in the field (Luck, Stewart, Simmons, & Rhemtulla, 2020; Rousselet et al., 2016; Rousselet & Pernet, 2011; Weissgerber, Garovic, Winham, Milic, & Prager, 2016; Weissgerber, Milic, Winham, & Garovic, 2015) but to date have been scantly reported in the literature for ERP effects of memory that traditionally just plot figures of bar graphs and ERP traces that inherently holding the potential to obscure the true nature of real patterns in the data (Rousselet et al., 2016; Weissgerber et al., 2016).

## 2.0 Method

The current investigation represented a re-analysis of data focused upon specific memory effects, which previously published in the domain of effects in metacognition (Muller, Sirianni, & Addante, 2020). Methods used in the participants, procedure, and analysis are re-stated in the sections below using similar text for clarity, and adapted where needed.

### 2.1 Participants

The total sample of participants consisted of 62 right-handed students free from neurological and memory problems from California State University, San Bernardino (CSUSB), as reported previously (Muller et al., 2020). Four participants’ data were not used due to noncompliance issues (pressed only 1 button throughout the task or ignored experimenter’s instructions) and one participant did not have usable data due to a experimenter error that resulted in the loss of that data. Two participants did not have usable EEG data due to excess motion artifacts/noise that resulted in a majority of trials being excluded from EEG and so were excluded from analyses. This presented a behavioral data set of N = 54 for the current study.

The majority of CSUSB participants were women (N = 48); 57% were Hispanic, 23% Caucasian, 11% Asian, and 10% identified as more than one ethnicity; for comparison, the prior study from UC Davis (Addante et al., 2012, N = 25) alternatively consisted of 24 % Hispanic, 54% Caucasian, and 25% Asian. The average age of CSUSB participants was 23.52 years old (*SD* = 4.82) and did not vary from that in the prior UC Davis cohort of 2012. None of the participants reported any visual, medical, or physical issues that would interfere with the experiment. Most participants spoke English as their first language (N = 47) and those whom who indicated speaking a different language first had been speaking English for an average of 16.73 years (*SD* = 4.74). Participants were recruited through a combination of methods including advertisements placed around CSUSB or through the school-wide research pool SONA. Participants recruited through advertisements were paid $10 an hour for sessions that lasted approximately three hours and participants recruited through SONA received 8 units of credit. Written informed consent was obtained for experimentation with human subjects and approved via the Institutional Review Board of California State University – San Bernardino, and data analyses observed the privacy rights of human subjects.

### 2.2 Procedure

The paradigm used was the same item- and source-memory confidence paradigm as that used in our prior reports (Addante et al, 2012a, 2012b, 2011, 2015; Roberts et al., 2018; Muller, Sirianni, & Addante, 2020), with slight modifications as noted below and in Muller et al (Figure 1). This paradigm consisted of an encoding phase containing four study sessions, in which participants studied 54 words in each session, and a retrieval phase, containing six test sessions in which the participant’s memory was tested for 54 words in each session. During test, participants viewed a total of 324 words, 216 of which were presented in the encoding phase and 116 of which were unstudied items (new items).

**Figure 1.**
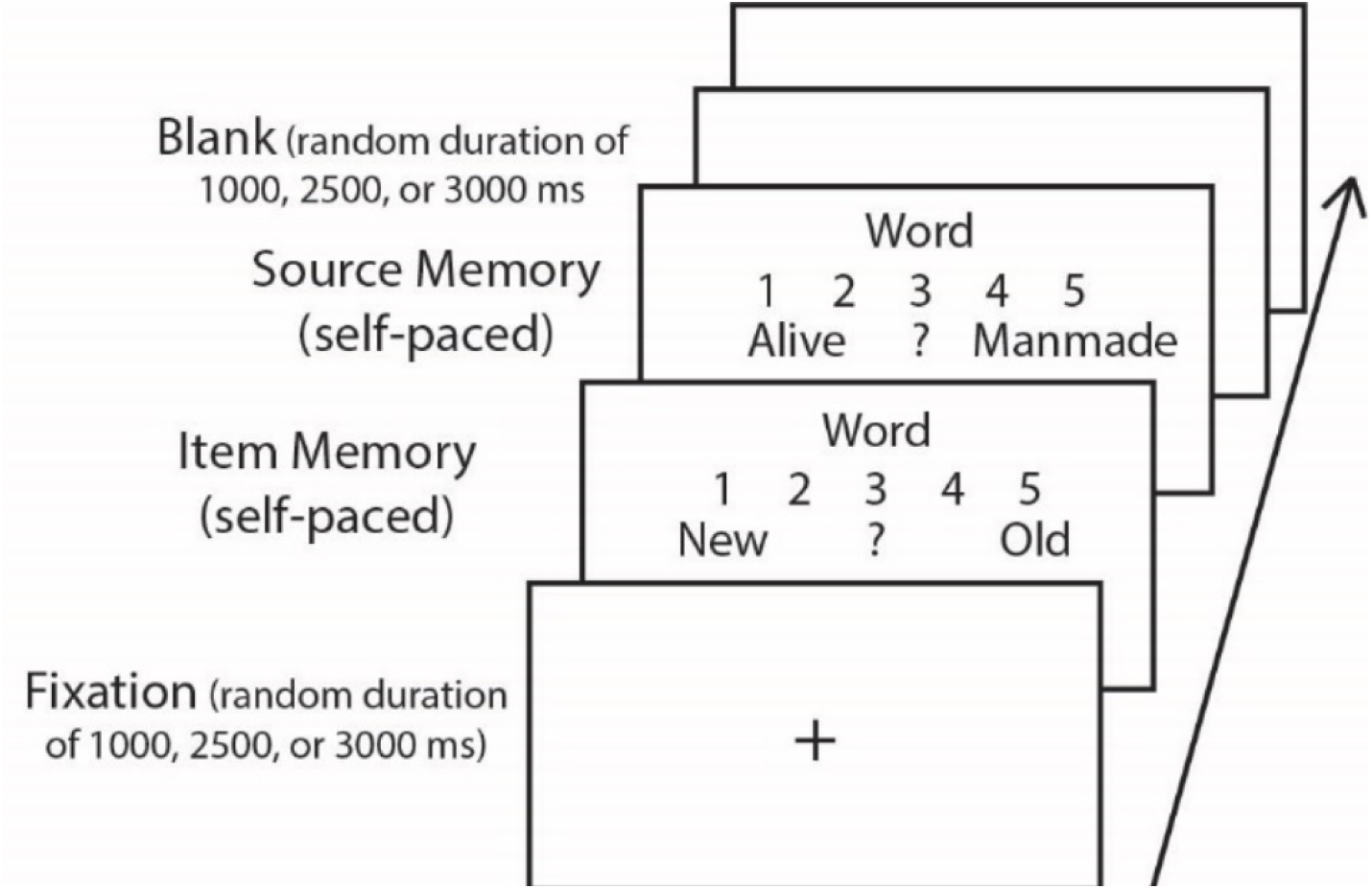
Retrieval paradigm. Memory test of item and source memory confidence ratings modified from Muller, Sirianni, & Addante (2020).

During the encoding phase, participants were given instructions to make a simple decision about the word presented (Figure 1). The participants were either asked to judge if the item was manmade or if the item was alive. The instructions were presented in one of two counterbalanced orders: ABBA or BAAB. The participants viewed four lists of 54 words during the encoding phase. The stimuli were presented on a black computer screen in white letters. To begin a trial, a screen with a small white cross at the center was presented for one of three randomly chosen inter-stimuli-interval (ISI) times: 1 second, 2.5 seconds, or 3 seconds. Then, the stimulus word appeared in the middle of the screen with ‘YES’ presented to the bottom left of the word and ‘NO’ presented to the bottom right of the word. The participants indicated their answer by pressing buttons corresponding to ‘yes’ and ‘no’ with their index and middle fingers, respectively. The response for this screen was self-paced by the participant. After the participants responded, they viewed a blank black screen at a random duration of 1 second, 2.5 seconds, or 3 seconds. After the blank screen, the small white cross appeared at the center of the screen to begin the next trial. This cycle continued until all 50 words in the all four lists were presented. Between each list, participants were read the instructions for the next task to ensure they correctly switched between the animacy and the manmade decision task. After the encoding phase was complete, the EEG cap was applied to the scalp of participants (see section below for details).

After the EEG cap was in place, the participant began the retrieval phase. The participants were read instructions asking them to judge if the stimulus word presented was old (studied during the encoding phase) or new (not studied before in the encoding phase) (Figure 2). As in the encoding phase, all stimuli words were presented in white font on a black screen. To begin a trial, a screen with a small white cross at the center was presented for one of three randomly chosen times: 1 second, 2.5 seconds, or 3 seconds. Then the participants were presented with a word in the middle of the screen, the numbers “1”, “2”, “3”, “4”, and “5” evenly spaced beneath the word, the word “New” on the left by the number “1”, and the word “Old” on the right under the number “5”. Participants pressed any number between “1” and “5” to indicate if they confidently believed the word was old (“5”), believe the word was old but was not confident (“4”), did not know if the word was old or new (“3”), believe the word was new but was not confident (“2”), or confidently believed the word was new (“1”). Participants were told to choose the response that gave us the most accurate reflection of their memory.

**Figure 2.**
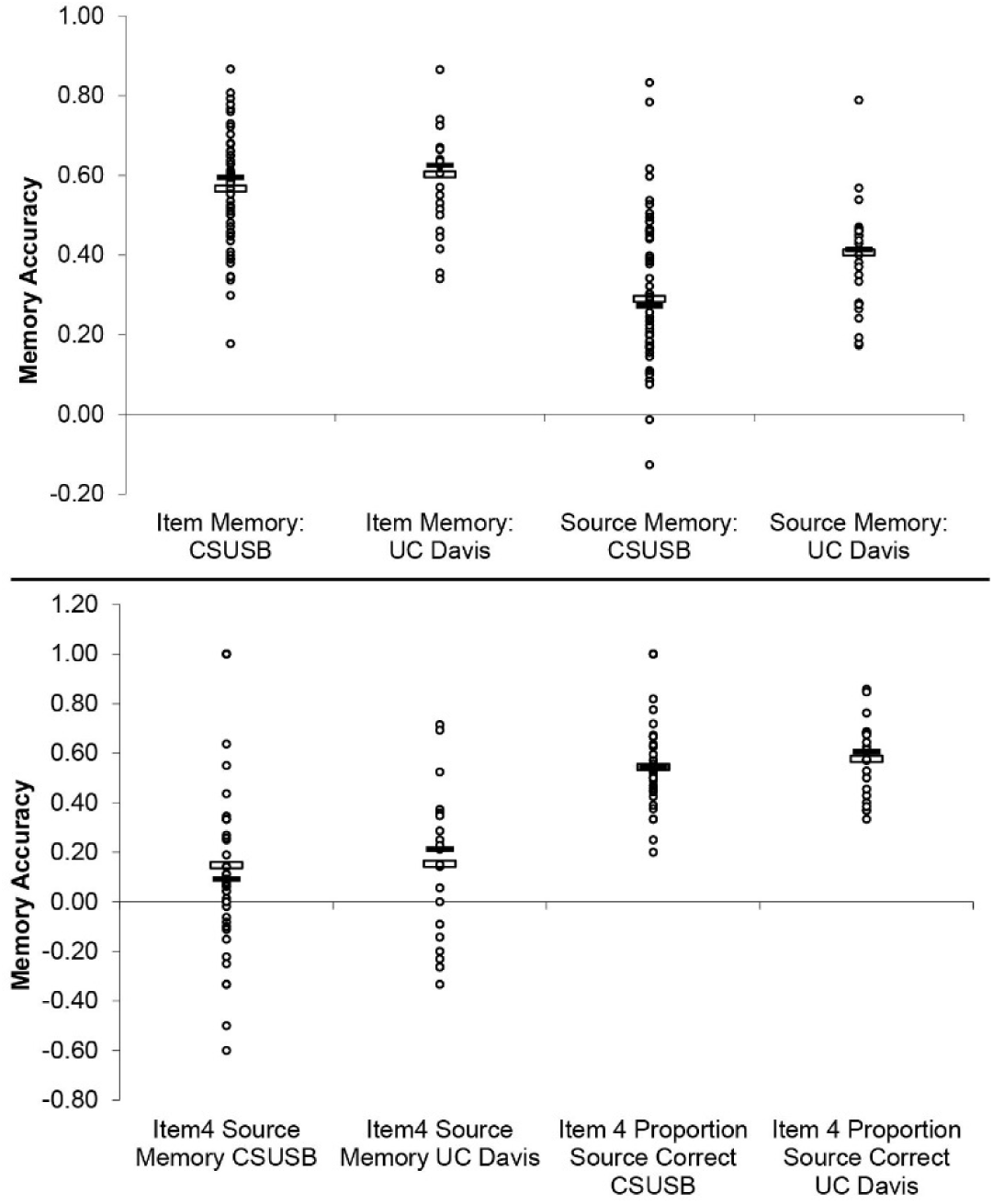
Memory performance measures for item recognition and source retrieval accuracy. Scatterplots indicate a data point for each participant’s data score for each measure denoted. Mean values for each condition are indicated by open white box, and median values are indicated by filled black box for each condition.

Immediately after the decision on item memory confidence, they were asked to answer if the word came from the animacy decision task or the manmade decision task. The word and numbers remained on the screen but this time, word “Alive” was presented on the left by the number “1”, and the word “Manmade” was presented on the right under the number “5”. Participants were told to choose the response that gave us the most accurate reflection of their memory and could respond that they confidently believed the word was from the animacy task (“1”), believe the word was from the animacy task but was not confident (“2”), did not know the source of the word or had replied in the question directly before that the word was new (“3”), believe the word was from the manmade task but was not confident (“4”), or confidently believed the word was from the manmade task (“5”). After that, a blank black screen was presented for a randomly chosen time of 1 second, 2.5 seconds, or 3 seconds (Figure 1). Participants were instructed to blink only during this blank screen and avoid blinking during the screens with a small cross or stimuli. The white cross was presented after the blank screen and the cycle continued until after the 10^th^ word has been presented. After the participant responded, the blank screen was presented and the next cycle of ten words were presented. Each session consisted of a list of 54 words. Six lists of 54 words were presented during the retrieval phase.

The retrieval phase of the paradigm also included a simple metacognitive question to participants for estimating their performance, which occurred once every ten trials and for which the data has been previously reported elsewhere for the metacognition effects (Muller, Sirianni, & Addante, 2020). The current investigation focused instead upon the memory related components of the study.

### 2.3 Electrophysiological Acquisition

Each subject was tested individually inside a private chamber. Stimulus presentation and behavioral response monitoring were controlled using Presentation software on a Windows PC. EEG was recorded using the actiCHamp EEG Recording System with a 32-channel electrode cap conforming to the standard International 10–20 System of electrode locations and was acquired at a rate of 1024 Hz. The EEG cap was sized while the participant’s face was wiped free of skin oil and/or makeup in preparation for attaching ocular electrodes. Five ocular electrodes were applied to the face to record electrooculograms (EOG): two above and below the left eye in line with the pupil to record electrical activity from vertical eye movements, two on each temple to record electrical activity from horizontal eye movements, and one electrode in the middle of the forehead in line horizontally with the electrode above the left eye as the ground electrode. EOG was monitored in the horizontal and vertical directions, and this data was used to eliminate trials contaminated by blinks, eye-movements, or other related artifacts. The EEG cap was placed on the participant’s head and prepared for electrical recording. Gel was applied to each cap site and impedances were lowered below 15 KOhms via gentle abrasion to allow the electrodes to obtain a clear electrical signal. Subjects were instructed to minimize jaw and muscle tension, eye movements, and blinking.

### 2.4 Electrophysiological Analyses

Physiological measurements of brain activity were recorded using EEG equipment from Brain Vision LLC. All EEG data was processed en masse using the ERPLAB toolbox using Matlab (Delorme and Makeig, 2004; Lopez-Calderon & Luck, 2014). The EEG data was first re-referenced to the average of the mastoid electrodes, passed through a high-pass filter at 0.1 hertz as a linear de-trend of drift components, and then downsampled to 256 hertz. The EEG data was epoched from 200 milliseconds prior to the onset of the stimulus to 1200 milliseconds after the stimulus was presented and then categorized based on performance group and response accuracy.

Independent components analysis (ICA) was performed using InfoMax techniques in EEGLab (Bell & Sejnowski, 1995) to accomplish artifact correction and then resulting data was individually inspected for artifacts, rejecting trials for eye blinks and other aberrant electrode activity. During ERP averaging, trials exceeding ERP amplitudes of +/- 250 mV were excluded. Using the ERPLAB toolbox (Lopez-Calderon & Luck, 2014), automatic artifact detection for epoched data was also used to identify trials exceeding specified voltages, in a series of sequential steps as noted below. Simple Voltage Threshold identified and removed any voltage below -100 ms. The Step-Like Artifact function identified and removed changes of voltage exceeding a specified voltage (100 uV in this case) within a specified window (200 ms), which are characteristic of blinks and saccades. The Moving Window Peak-to-Peak function is commonly used to identify blinks by finding the difference in amplitude between the most negative and most positive points in the defined window (200 ms) and compared the difference to a specified criterion (100 uV). The Blocking and Flatline function identified periods in which the voltage does not change amplitude within the time window. An automatic blink analysis, Blink Rejection (alpha version), used a normalized cross-covariance threshold of 0.7 and a blink width of 400 ms to identify and remove blinks (Luck, 2014). ERPs of individual subjects were combined to create a grand average, and mean amplitudes were extracted for statistical analyses, and a 30 Hz low pass filter was applied to averaged ERPs. During ERP averaging, trials exceeding ERP amplitudes of +/- 250 mV were excluded. Data is accessible upon request without restriction, as is any code used to analyze the data- which for the current study involved GUI-based operations in ERPLab. Authors encourage communications from those interested in innovative additional analyses.

In order to maintain sufficient signal-to-noise ratio (SNR), all comparisons relied upon including only those subjects whom met a criterion of having a minimum number of 12 artifact-free ERP trials per condition being contrasted (Addante, Ranganath, & Yonelinas, 2012; Gruber and Otten, 2010; Kim et al., 2009; Otten et al., 2006; c.f. Luck 2016). Topographic maps of scalp activity for differences between conditions were created to assess the spatial distribution of the observed effects. Because the nature of the current investigation was based upon clear a priori-defined hypotheses derived from prior findings (Addante et al., 2012), analyses thus utilized planned paired t-tests to assess differences between the targeted conditions, latencies, and scalp locations noted. In cases where exploratory analyses were conducted to explore unplanned comparisons, ANOVA was used to qualify potential differences that may exist. Participants were excluded from any particular analyses if they did not have responses in both/all of the comparisons for a given contrast.

## 3.0 Results

### 3.1 Behavioral Results

#### 3.1.1 Item Recognition Memory Performance

Recognition memory response distributions for recognition of old and new items are displayed in Table 1 and Figure 2 (adapted from Muller et al., 2020). Item recognition accuracy was calculated as the proportion of hits (*M* = .81, *SD* = .11) minus proportion of false alarms (*M* = .24, *SD* = .14) (i.e. pHit-pFA). Participants performed item recognition at relatively high levels (*M* = .57, *SD* = .14, SE = .02) which was greater than chance (i.e. zero), *t*(53) = 29.36, *p* < .001. In addition, participants’ accuracy for high confidence item recognition trials (‘5’s’: M = .82, SD = .12) was significantly greater than low confidence item recognition trials (‘4’s’: M = .39, SD = .38), *t*(53) = 8.78, *p* < .001 (Figure 2).

**Table 1.**
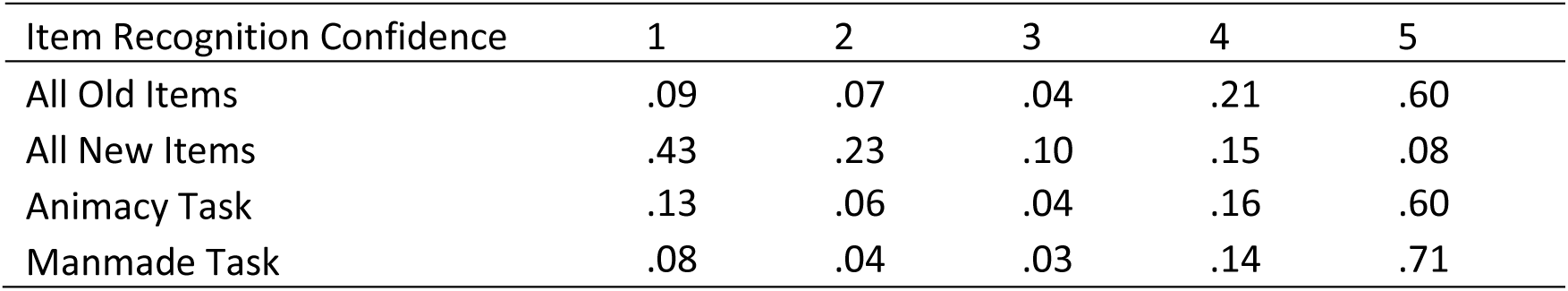
Distribution of Responses for Each Item Response as a Proportion of All Memory Responses

#### 3.1.2 Source Memory Performance

Source memory response distributions for old and new items are displayed in Table 2 and Figure 2 (adapted from Muller et al., 2020). General source memory accuracy values were collapsed to include high- and low- confidence source responses which were then divided by the sum of items receiving a correct and incorrect source response, to calculate the proportion as was done in prior reports (R. J. Addante, Ranganath, Olichney, & Yonelinas, 2012; R. J. Addante, Ranganath, & Yonelinas, 2012; R. J. Addante et al., 2011; Muller et al., 2020; Roberts, Clarke, Addante, & Ranganath, 2018). Mean accuracy for general source memory was .31 (*SD* = .19, SE = .03) and was reliably greater than chance, *t*(55) = 11.78, *p* < .001.

**Table 2.** Distribution of Responses for Each Source Response as a Proportion of All Memory Responses

| Source Recognition Confidence | 1 | 2 | 3 | 4 | 5 |
| --- | --- | --- | --- | --- | --- |
| All Old Items | .14 | .14 | .22 | .17 | .33 |
| All New Items | .05 | .08 | .70 | .09 | .08 |
| Animacy Task | .24 | .17 | .27 | .16 | .16 |
| Manmade Task | .11 | .11 | .17 | .2 | .41 |

#### 3.1.3 Between-Experiment Comparisons

Because the current study sought to replicate the neurophysiological findings from a prior study (R. J. Addante, Ranganath, & Yonelinas, 2012) and extend it in novel ways, we also sought to first assess the extent to which the two experiments may or may not differ in their underlying performance for item and source memory, using two-tailed between-group t-tests. In the most general test of performance, item recognition accuracy (proportion of hits – proportion of false alarms), the group in the current study (N = 54, M = .57, SD = .14, SE = .02, ) and the group from the former study at UC Davis (N = 25 M = .60, SD = .14, SE = .03) were found not to differ, t(77) = .976, p = .332 (Figure 2). In general measures of source memory (proportion of source correct minus proportion of misattributed source), the group of the current study at CSUSB exhibited worse source memory overall (M = .31, SD = .19, SE = .03) than did the UC Davis group of 2012 (M = .410, SD = .16, SE = .03), t(77) = 2.20, p = .031, though as noted above, still performed at above-chance levels, indicating their successful source discriminability in memory.

#### 3.1.4 Item and Source Memory Combination performance

The current study was adapted from prior work that reported uniquely different response profiles for correct source judgements that were preceded by high and low levels of item recognition confidence hits (R. J. Addante, Ranganath, & Yonelinas, 2012). The same analysis was performed on the current data. The memory response combination of low-confidence item recognition hits (item4) + correct source memory was provided at the same proportion for both experiments (CSUSB: M = .572, SD = .18, SE = .03; UC Davis: M = .576, SD = .15, SE = .03), and which did not differ reliably t(71) = .096, p = .923 (Figure 2). In each of the studies (at CSUSB and past at UC Davis) source memory for the low-confidence hits (item4) was performed at above-chance levels of discriminability (CSUSB: t(47) = 2.85, p = .006; UC Davis: t(24) = 2.50, p = .019).

More broadly, the broad effect of differential general source memory performance was further explored more specifically for its constituent sub-conditions of accuracy of the high- (item5) and low-(item4) confidence levels of item recognition among each experiment’s groups, respectively. For high confident (item5) hits, experiments were found to differ, (t(77) = 2.39, p = .019; CSUSB M = .36, SD = .20, SE = .03; UC Davis M = .47, SD = .16, SE = .03), but for the key contrast of interest to the current study (low confidence hits [item4]) the two experiments were not found to differ in source memory accuracy, t(71) = .096, p = .923 (CSUSB M = .145, SD = .35, SE = .05; UC Davis M = .152, SD = .31, SE = .06) (Figure 2). Thus, overall there were no differences observed between experiments for the main analysis aim of the current study (source memory accuracy of low-confidence item hits), but the only differences evident were for source memory accuracy of the high-confidence hits (item5 responses), which were apparently driving the overall differences observed earlier between experiments in source memory accuracy. Importantly, the current experiment was found to be functionally the same, in both task and performance, as the preceding experiment from the prior one that motivated it (Addante et al., 2012), including measures of item recognition accuracy and the specific source memory conditions of interest.

#### 3.1.5 Response Speed for Recognition Memory Judgments

Reaction times for each item response are shown in Table 3 while reaction times for each source response are shown in Table 4 (adapted from Muller et al., 2020) (see also Figure 3). In item recognition judgments, participants responded significantly faster when identifying hits (M = 2280 ms, SD = 549 ms, SE = 74.84 ms) than misses (M = 2982 ms, SD = 923 ms, SE = 125.68 ms), *t*(53) = -7.04, *p* < .001, false alarms (M = 2762 ms, SD = 850 ms, SE = 110 ms), *t*(53) = -4.03, *p* < .001, and correct rejections (M = 2626 ms, SD = 799 ms, SE = 108.33 ms), *t*(53) = -4.09, *p* < .001. They also responded significantly faster when identifying correct rejections than misses, *t*(53) = 3.30, *p* = .001, and misses to false alarms, *t*(53) = 2.04, *p* = .03. There were no significant differences between the reaction times for false alarms and correct rejections, *t*(53) = 0.93, *p* = .35. In addition, participants responded significantly faster to high confidence item recognition trials (5’s’) (*M* = 1897 ms, *SD* = 397 ms) than low confidence item recognition trials (4’s’) (*M* = 2834 ms, *SD* = 1032 ms), *t*(49) = -8.10, *p* < .001.

**Figure 3.**
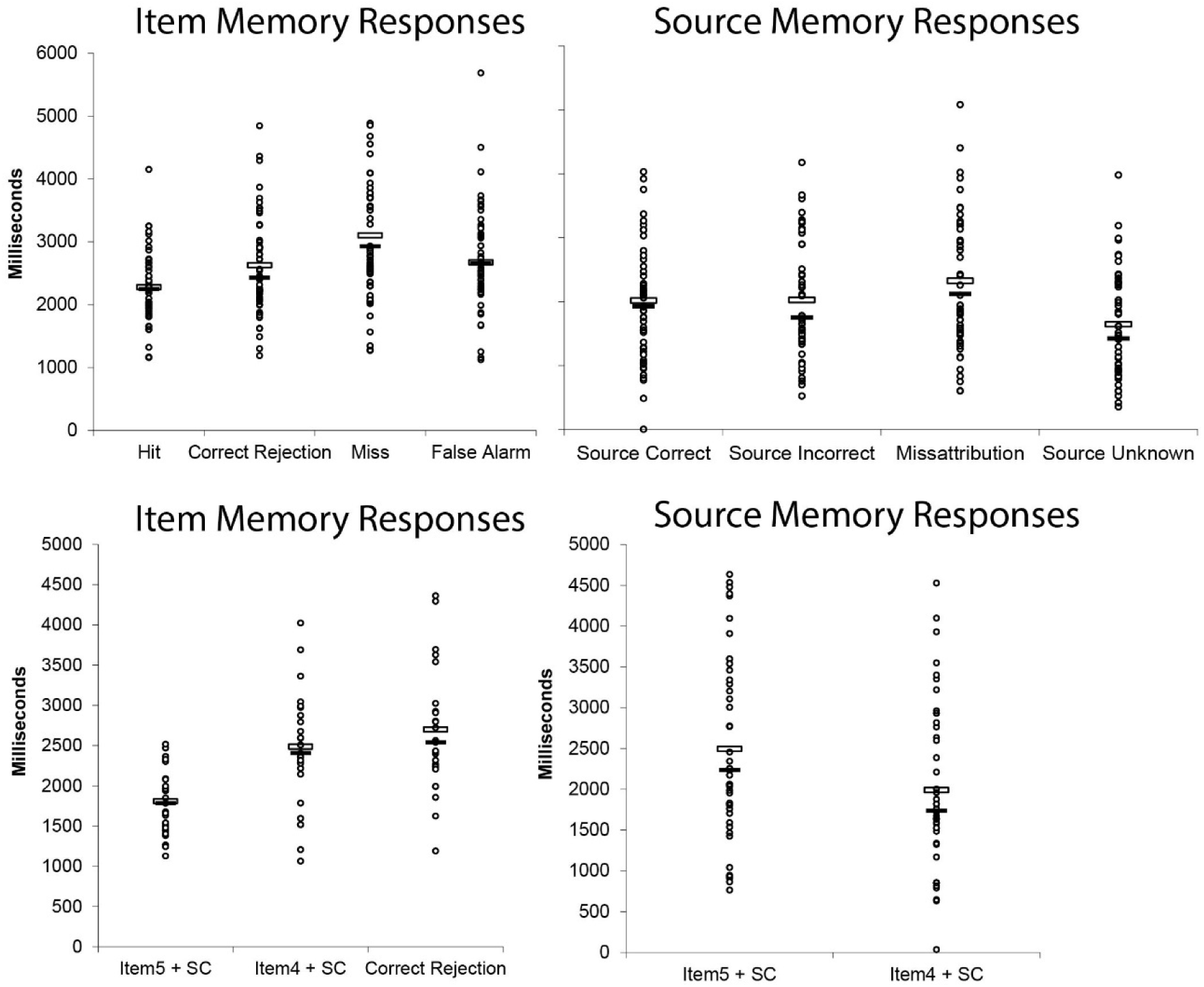
Response times for item recognition and source memory judgments. Scatterplots indicate a data point for each participant’s data score for each measure denoted. Mean values for each condition is indicated by open white box, and median values are indicated by filled black box for each condition.

**Table 3.** Average Reaction Times for Each Item Memory Response

| Item Reaction Times | 1 | 2 | 3 | 4 | 5 |
| --- | --- | --- | --- | --- | --- |
| All Old Items | 2547<br>(1067) | 3295<br>(1534) | 3151<br>(1678) | 2682 (752) | 1852 (394) |
| All New Items | 2205 (671) | 3014<br>(1200) | 2897<br>(1858) | 2999 (868) | 2085 (820) |
| Animacy Task | 2451 (990) | 3355<br>(1128) | 3376<br>(1352) | 2732 (976) | 1853 (378) |
| Manmade Task | 2470 (782) | 3425<br>(1369) | 3254<br>(1391) | 2947<br>(1174) | 1765 (339) |

*Note.* Values are in milliseconds with standard deviations in parentheses.

**Table 4.**
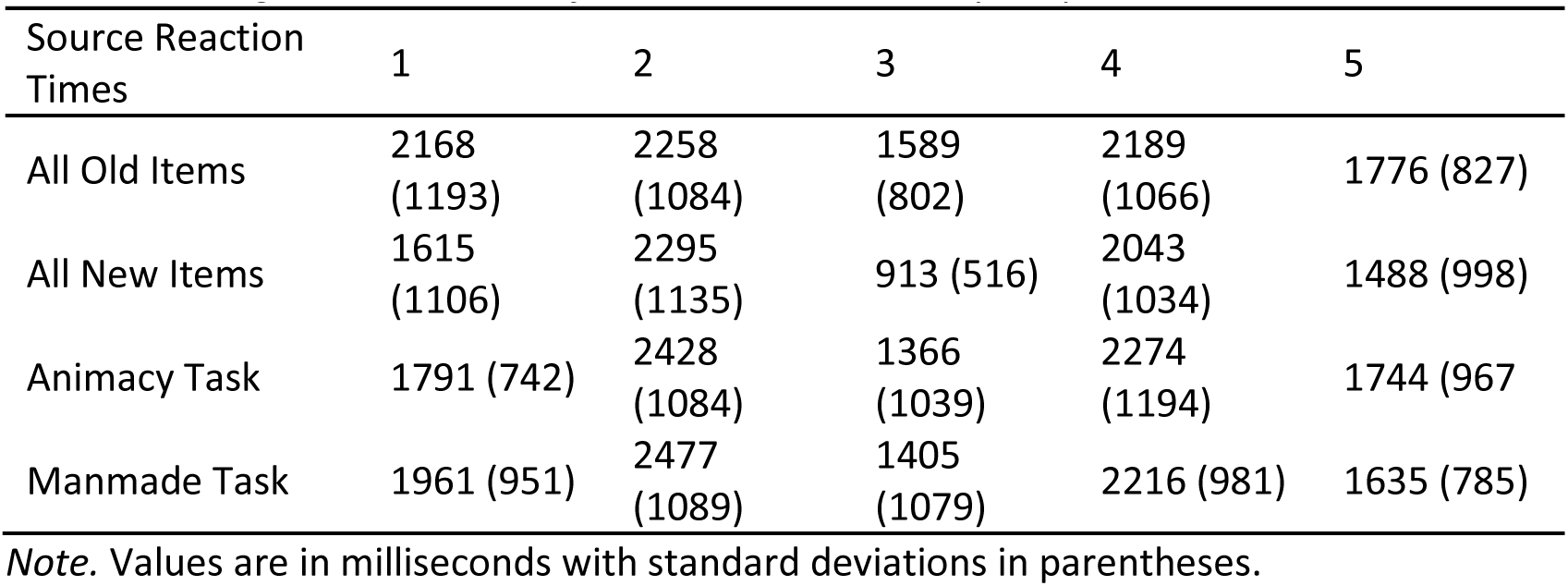
Average Reaction Times for Each Source Memory Response

| Source Reaction Times | 1 | 2 | 3 | 4 | 5 |
| --- | --- | --- | --- | --- | --- |
| All Old Items | 2168<br>(1193) | 2258<br>(1084) | 1589<br>(802) | 2189<br>(1066) | 1776 (827) |
| All New Items | 1615<br>(1106) | 2295<br>(1135) | 913 (516) | 2043<br>(1034) | 1488 (998) |
| Animacy Task | 1791 (742) | 2428<br>(1084) | 1366<br>(1039) | 2274<br>(1194) | 1744 (967) |
| Manmade Task | 1961 (951) | 2477<br>(1089) | 1405<br>(1079) | 2216 (981) | 1635 (785) |

*Note.* Values are in milliseconds with standard deviations in parentheses.

The finding of item judgment response time differences for memory confidence persisted even when source memory was held constant, as comparing the low confidence item judgments (‘4’s’) that had source-correct judgments (*M* = 2767 ms, *SD* = 1077 ms) to high-confidence item judgments (‘5’s’) that had source-correct judgments (*M* = 1935 ms, *SD* = 618 ms) found that the high confidence hits were still reliably faster (*t*(47) = 6.96, *p* < .001)^1^.

#### 3.1.6 Response Speed for Source Memory Judgments

Reaction times were also assessed for the source memory judgments given by participants during the test (immediately following each item memory judgment). First, we analyzed the general source memory conditions of times when participants remembered the source correctly, as well as times when people provided the incorrect source (defined as instances when people provided the wrong source misattribution or said the source was unknown, e.g. Addante et a., 2011; 2012a). There were no reliable differences in response speed (in milliseconds) for source correct (M = 2016, 856, SE = 117) and source incorrect (M = 2026, SD = 886, SE = 121) conditions, t(52) = -.127, p = .90 (Figure 3). When the source incorrect condition was assessed separately for its constituent parts, it revealed that the ‘source unknown’ responses (M = 1614, SD = 810, SE = 111) were faster than both correct and misattribution (M = 2232, SD = 1024, SE = 140) of source memories (t(53) = 3.96, p <.001, t(53) = 5.80, p < .001), respectively), and the misattributions were also slower than correct source memory responses (t(53) = 2.63, p = .011).

When the specified conditions of item confidence + source judgments were directly assessed for the reaction speed of the source memory responses, it revealed that high confident recognition hits (item5) + source correct trials were responded to significantly slower than low confident recognition hits (item4) + source correct trials (t(45) = 5.05, p < .001) (Figure 3). This represented the opposite pattern that had been observed in the item memory response times for these conditions, when the item5+ source correct responses were instead found to be faster than the item4+source correct trials (see Item RT section noted above). In order to qualify these patterns of findings for the item and source memory response times across the item and source memory judgments, respectively, these conditions were formally subjected to a 2×2 ANOVA with factors of condition (item5+source correct, item4+source correct) and the judgment response time (item memory, source memory), which identified a significant condition by response time interaction, F(1,180) = 27.53, p < .001. These results indicate that for these two particular conditions of correct source memory, people responded faster for the high confident recognition hits but slower for their subsequent source memory judgments, whereas the opposite pattern was true for the low confident recognition hits-which were slower during recognition but faster during source memory retrieval.

### 3.2 Electrophysiological Results

#### 3.2.1 Recognition Memory ERPs

The approach of the analyses was designed to parallel those of the study which motived this investigation (R. J. Addante, Ranganath, & Yonelinas, 2012), and thus follow the same step-wise progression from general to specific memory measures, while extending them with additional measures of variance that were unavailable in prior ERP studies of memory. First, general recognition memory was analyzed by comparing ERPs for correctly identified old items (hits: combined responses of ‘4’ and ‘5’) to correctly identified new items (correct rejections: combined responses of ‘1’ and ‘2’) (this basic analysis of the current dataset was previously reported in Muller et al, 2020). We analyzed the FN400 effect at the same site as reported in Addante et al. (2012a) (Cz) and it was found to be a significant effect of hits being more positive-going than correct rejections from 400 to 600ms (*t*(53) = 3.76, *p* < .001). Consistent with prior findings, this FN400 effect was then found to then shift towards the left parietal region during later latencies of 600-800ms, with ERPs for hits becoming significantly more positive than correct rejections at left parietal site P3, (t(53) = 2.97, p = .004, demonstrating the canonical LPC effect (Figure 4).

**Figure 4.**
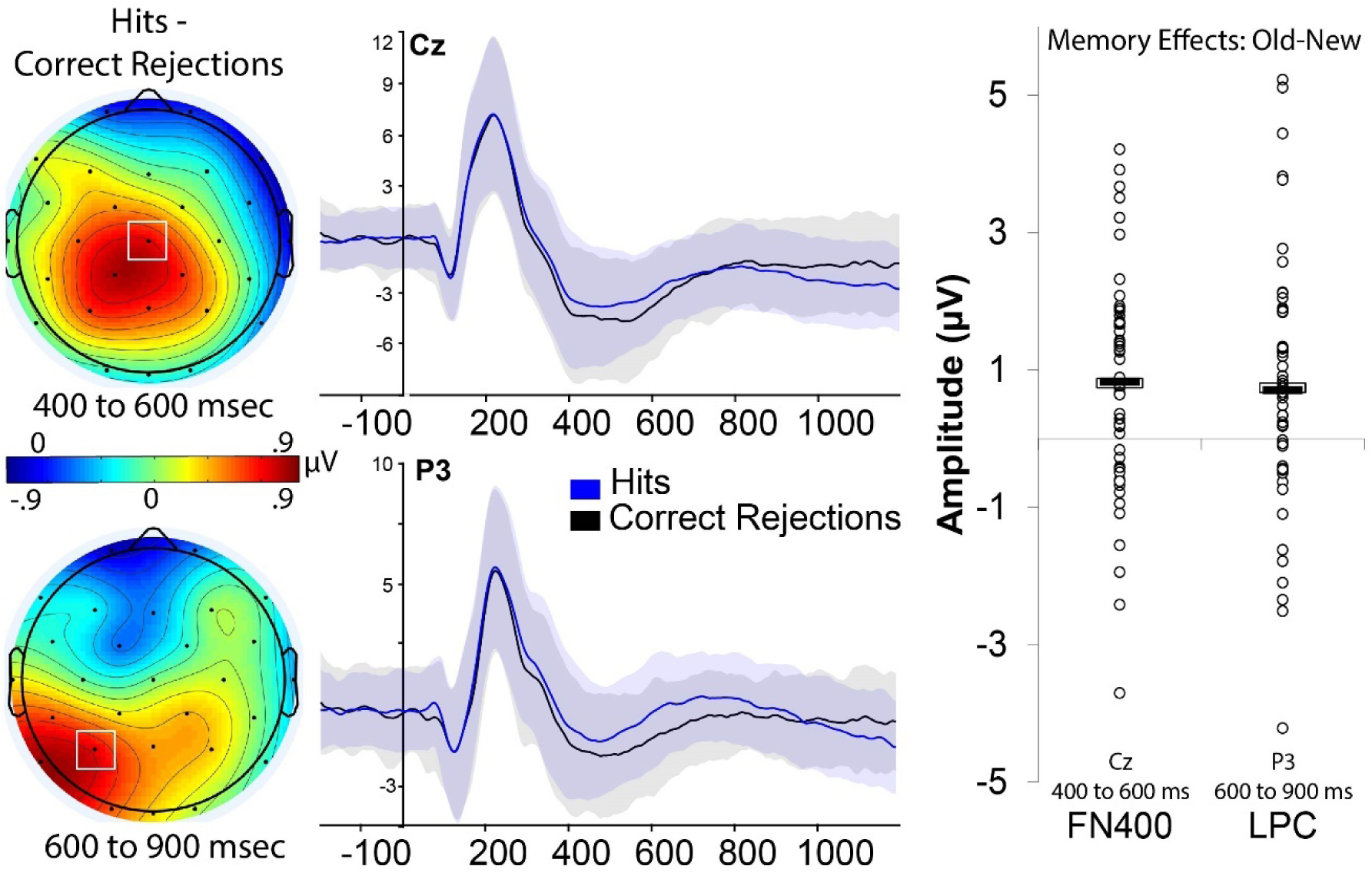
ERPs for basic recognition memory. Topographic maps indicate the difference waves for ERPs of hits minus corrections for each time latency noted, and the magnitude of any differences corresponding the values shown on the color scale indicated, with warmer colors reflecting more positive differences in voltage. ERPs show milliseconds on the x-axis and μV on the y-axis, with measures of variance (standard error of the mean) indicated by the background shading in the color of each condition noted. Scatterplots indicate a data point for each participant’s data score for each measure of the old-new difference (hits minus correct rejections); mean values for each condition is indicated by open white box, and median values are indicated by filled black box for each condition.

#### 3.2.2. Item recognition Confidence ERPs

Second, we more-specifically investigated differences in ERPs for high- and low-confidence hits, to evaluate the consistency with, and replicability of, related findings reported in neuropsychological patients (Addante et al., 2012b). ERPs for high confidence hits (response ‘5’) were more positive-going than ERPs for low-confidence hits (item response of ‘4’), revealing the same pattern of FN400 effects at mid-frontal sites (Fc1) from 400-600 ms (*t*(33) = 2.45, *p* = .019) and LPC effects at left parietal site (P3) from 600-900ms as was reported among the prior study of hippocampal amnesia patients and controls, respectively, by Addante et al., (2012b) (*t*(33) = 2.91, *p* = .006), and extending it with the additional measures of variance obtained from our relatively large sample size of the current study (N=54) than were available in the smaller sample sizes of clinical patients tested previously (N=3, N=6). Similar significant effects were also found at adjacent sites (i.e.: Fz, Cp5) and latencies (i.e.: 300-500ms, 700-900ms), consistent with the variable ranges typically observed across the literature (Figure 5).

**Figure 5.**
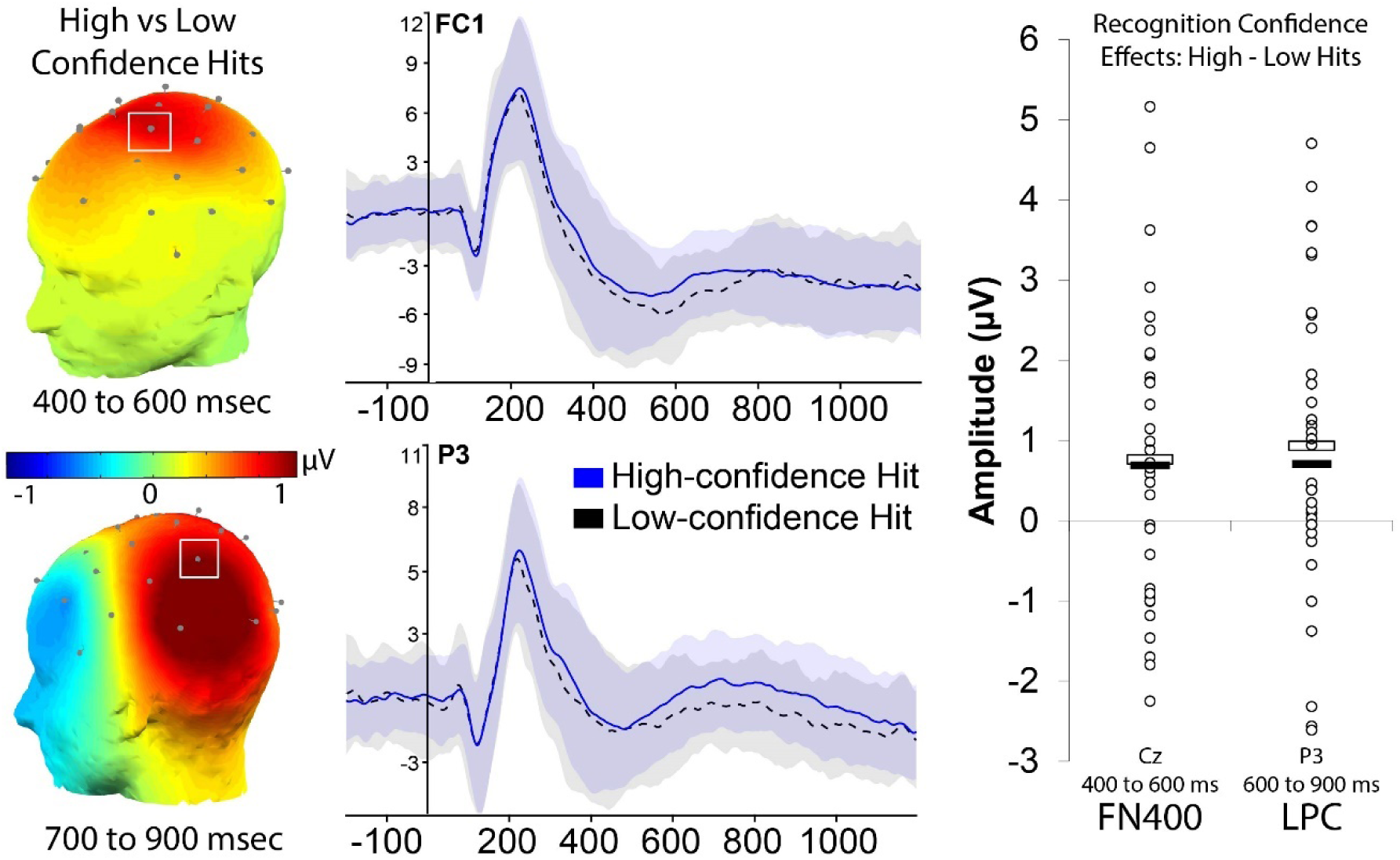
ERPs for specific recognition memory confidence levels. Topographic maps indicate the difference waves for ERPs of high confidence hits minus low confidence hits for each time latency noted, and the magnitude of any differences corresponding the values shown on the color scale indicated, with warmer colors reflecting more positive differences in voltage. ERPs show milliseconds on the x-axis and μV on the y-axis, with measures of variance (standard error of the mean) indicated by the background shading in the color of each condition noted. Scatterplots indicate a data point for each participant’s data score for each measure of the difference between high confidence hits minus low confidence hits; mean values for each condition is indicated by open white box, and median values are indicated by filled black box for each condition.

#### 3.3.3 Source Memory ERPs

Third, ERPs for source memory were analyzed by comparing judgments of each correct and incorrect source memory responses to correct rejections. For source correct judgments, an FN400 effect was evident from 400-600 ms at central site Cz, again replicating findings from prior studies (Addante et al., 2012), *t*(53) = 3.58, *p* < .001. During later latencies of 600-900 ms, correct source judgments elicited the canonical LPC effect of recollection (R. J. Addante, Ranganath, Olichney, et al., 2012; R. J. Addante, Ranganath, & Yonelinas, 2012; Friedman, 2013; M. D. Rugg & Curran, 2007) maximal over left parietal site P3, *t*(53) = 3.37, *p* = .001. For source incorrect judgements, an FN400 effect was similarly evident from 400-600 ms at Cz (*t*(53) = 2.86, *p* = .006), but during the later latencies of 600-900 ms for the LPC source incorrect ERPs were not significantly different than correct rejections (*t*(53) = 1.87, *p* = .067) (Figure 6).

**Figure 6.**
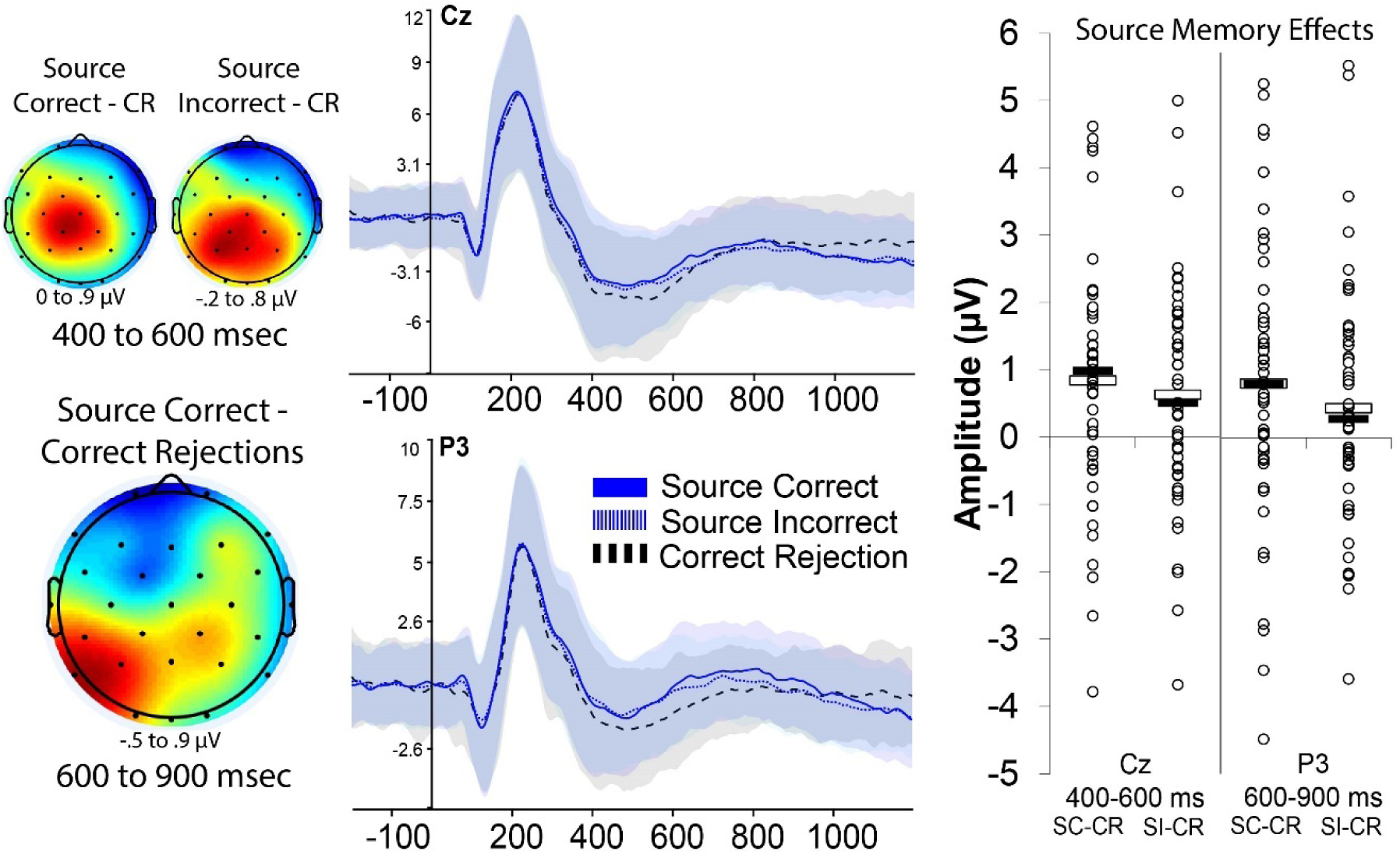
ERPs for source memory. Topographic maps indicate the difference waves for ERPs of source memory conditions minus correct rejections for each time latency noted, and the magnitude of any differences corresponding the values shown on the color scale indicated, with warmer colors reflecting more positive differences in voltage. ERPs show milliseconds on the x-axis and μV on the y-axis, with measures of variance (standard error of the mean) indicated by the background shading in the color of each condition noted. Scatterplots indicate a data point for each participant’s data score for each measure of the difference between source memory conditions and correct rejections; mean values for each condition is indicated by open white box, and median values are indicated by filled black box for each condition.

#### 3.3.4 ERPs for Item and Source Memory Combinations

The original report of combinations of item and source memory confidence ERPs (Addante et al., 2012) reported that it was associated with a late negative-going ERP effect when compared to correct rejections, which was predominantly occurring at broad frontal-central sites from 800-1200ms. Thus, we pursued the same analytic approach here, and also included the 400-600ms and 600-900ms latencies to verify if the condition exhibited any association with traditional effects of item familiarity (FN400) or recollection (LPC) (it did not in previous studies).

We assessed ERPs for instances of low confidence recognition hits (‘4’s’) with source-correct memories, as compared to correct rejections, and found evidence of the same negative-going effect from 800-1200 ms occurring at left frontal-central electrode sites that had been previously reported in Addante et al., (2012a, 2012b) (Fc1: t(26) = -2.42, p = .022; Cz: t(26) = - 2.52, p = .018) as well as more widespread regions of central-parietal sites as well, thereby replicating the prior findings with a larger sample that doubled in size (from an N=13 to an N=27) (Figure 7)^2^. Analyses also confirmed prior reports that there was no LPC effect evident for this condition at left parietal sites such as P3 from 600-900 ms (t(26) = 0.16, p = .873), despite there being accurate source memory retrieval, nor was there any evidence for a traditional FN400 effect of item familiarity from 400-600ms (t(26) = -0.06, p = .955), despite there being above-chance item recognition success (Figure 2).

**Figure 7.**
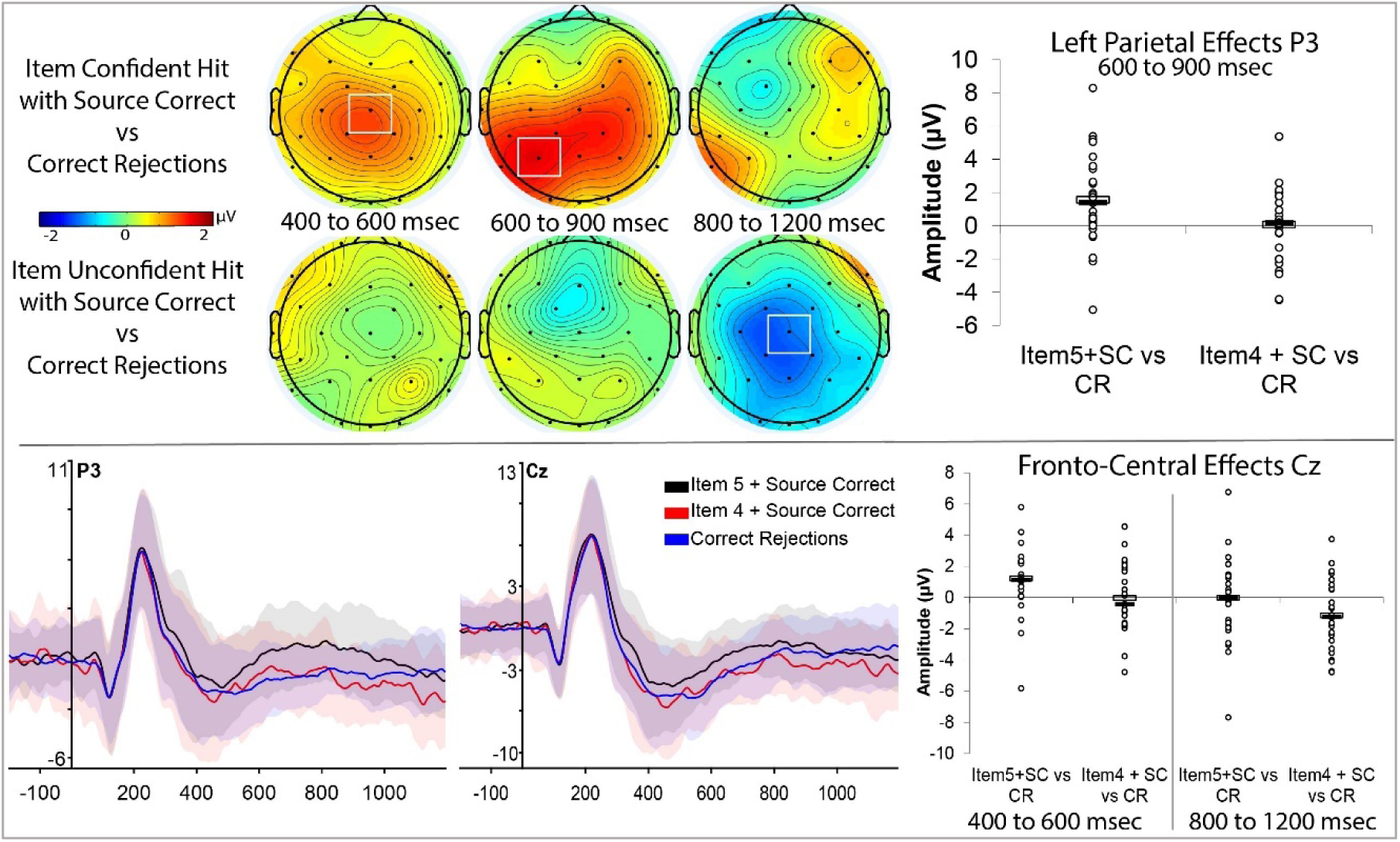
ERPs for item and source memory combination conditions. Topographic maps indicate the difference waves for ERPs of item+source memory conditions minus correct rejections for each time latency noted, and the magnitude of any differences corresponding the values shown on the color scale indicated, with warmer colors reflecting more positive differences in voltage. ERPs show milliseconds on the x-axis and μV on the y-axis, with measures of variance (standard error of the mean) indicated by the background shading in the color of each condition noted. Scatterplots indicate a data point for each participant’s data score for each measure of the difference between the item+source memory conditions and correct rejections; mean values for each condition is indicated by open white box, and median values are indicated by filled black box for each condition.

In contrast, ERPs for high confidence item hits (‘5’s’) with correct source memory did elicit an LPC when compared to correct rejections at left parietal site of P3 from 600-900ms, *t*(26) = 2.98, *p* = .006 (Figure 7), which visual inspection of the data revealed as a widespread bilateral parietal effect, and which replicated prior findings for recollection-related processing of high confident recognition and accurate source memory (R. J. Addante, Ranganath, & Yonelinas, 2012; Wilding & Rugg, 1996; Woodruff, Hayama, & Rugg, 2006; Yu & Rugg, 2010). Accordingly, high confident recognition hits with correct source memory also exhibited an earlier FN400 effect at Cz, (t(26) = 2.82, p = .008), also consistent with similar prior findings.

#### 3.3.5 Exploratory analysis of negative-going central memory ERP effects

The preceding findings indicated a fronto-central/parietal ERP effect for memory of episodic context by successful low-confidence recognition hits, occurring from approximately 800-1200 msec post-stimulus onset. These effects are consistent with prior reports using the same paradigm (Addante et al., 2012a, 2012b) where they were similarly characterized in normative and neuropsychological patients. The functional significance of this otherwise-novel ERP effect of memory remains relatively unclear, so to be thorough, we conducted an exploratory follow-up analysis that investigated if similar effects could be observed in related contrasts of mnemonic conditions in which episodic context was retrieved. This analysis was exploratory, and unplanned, and thus was based upon visual inspection of the ERPs from 800-1200ms in the preceding analyses including comparisons of hits vs correction rejections, source correct vs correct rejections, source incorrect vs correct rejections. Each contrast was found to exhibit a significant effect at electrode site Cz from 800 to 1000 msec: hits (t(53) = -2.95, p = .004), source correct (t(53) = -2.32, p = .024), and source incorrect (t(53) = -2.85, p = .006) (Figure 8). In order to ascertain if these observed effects may all represent a form of the same ERP effect, each memory contrast’s difference wave was subsequently included in an exploratory 4×1 ANOVA that also included the main negative-going ERP effect of item4+source correct vs correct rejections as well, and the memory effects were found not to vary (F(3,185) = .38, p = 767).

**Figure 8:**
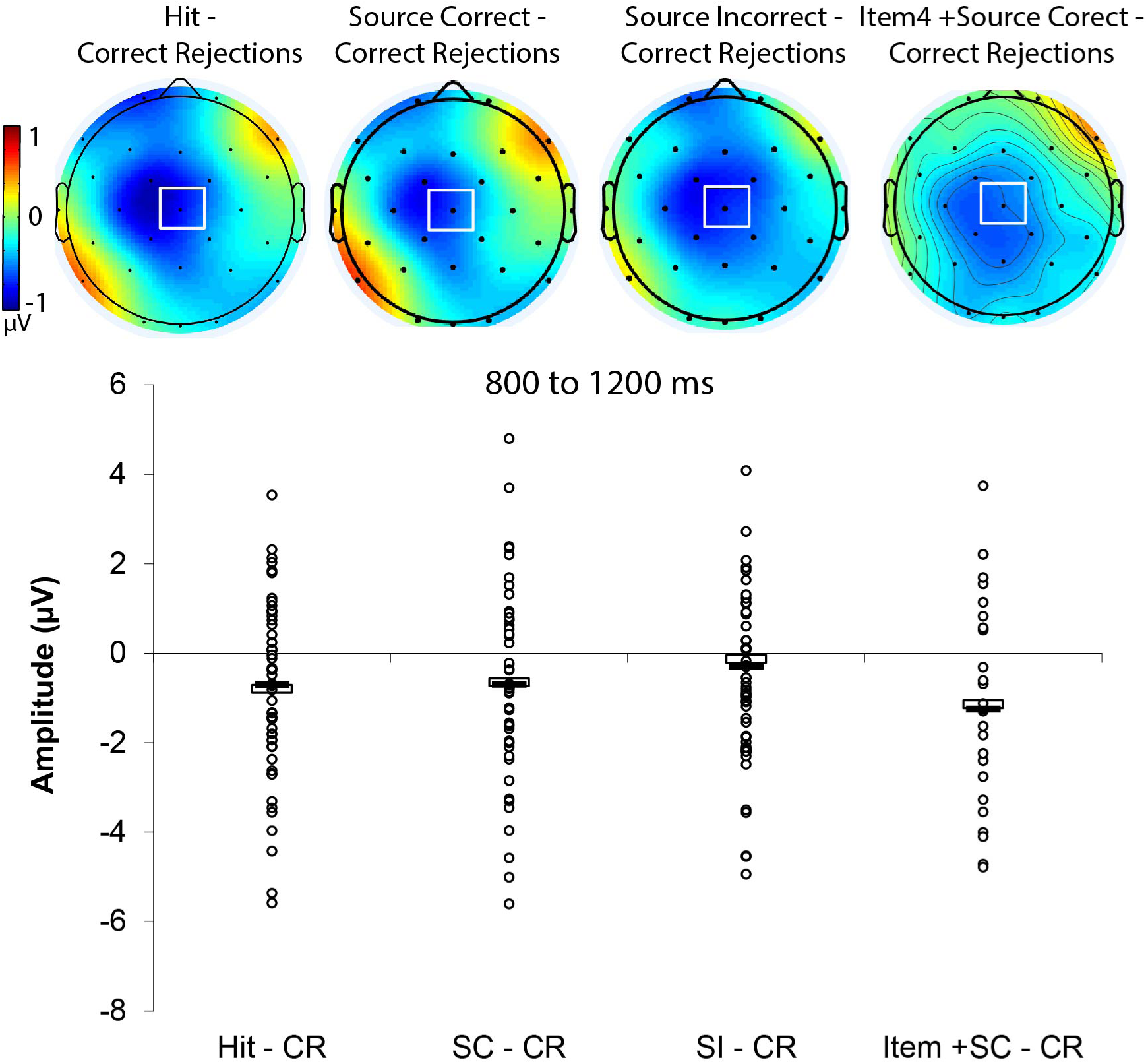
Late Central Negativity (LCN) Effect. Topographic maps indicate the difference waves for ERPs of source memory conditions minus correct rejections for each time latency noted, and the magnitude of any differences corresponding the values shown on the color scale indicated with warmer colors reflecting more positive differences in voltage. Scatterplots indicate a data point for each participant’s data score for each measure of the difference between source memory conditions and correct rejections; mean values for each condition is indicated by open white box, and median values are indicated by filled black box for each condition.

## 4.0 Discussion

### 4.1 General Summary of Findings

By using a well-established memory paradigm of item and source memory confidence measures with ERPs, we were able to replicate several established memory effects from the literature, and added to those findings with novel measures of variance characterizing the broader nature of these effects that has been missing from the literature thus far. We first identified basic, canonical effects of memory: that old items are remembered better, faster, and with more positive voltage than new items (Tables 1 and 2) (Sanquist, Rohrbaugh, Syndulko, & Lindsley, 1980), and furthermore that ERPs for these items were associated with the canonical effects of the FN400 and LPC that are traditionally viewed as the putative neural correlates of familiarity- and recollection-based memory processing (Figures 8 and 9) (R. J. Addante, Ranganath, & Yonelinas, 2012; Curran, 2000; Friedman, 2013; P. A. Leynes, Bruett, Krizan, & Veloso, 2017; Mecklinger & Bader, 2020; Mecklinger, 2006, 2010; Muller et al., 2020; M. D. Rugg & Curran, 2007).

In item recognition confidence measures, performance was better and faster for the high confidence hits than the low-confidence hits, and ERPs revealed the same FN400 and LPC effects as had been found in prior studies of neuropsychological patients (R. J. Addante, Ranganath, Olichney, et al., 2012). We also identified behavioral and physiological effects for source memory, revealing that an FN400 effect was evident for both the conditions of source correct and source incorrect trials, but that the LPC was evident only for the source correct trials (Figure 10), consistent with earlier findings from this paradigm (Addante et al., 2012a, b) and consistent with theoretical models positing recollection and familiarity as dual processes of episodic memory (Yonelinas, 2002; 2004; 2010; Eichenbaum et al., 2007; see also ERP review of Rugg & Curran, 2007).

The main finding of the current study was that it was also able to extend previously-reported ERP effects of memory combinations for item and source retrieval that have remained relatively unexplored in the field. In particular, Addante et al. (2012a) reported a novel late fronto-central negative-going ERP effect for instances in which participants provided low-confidence item memory hits that still had accurate source memory judgments. The current study provided a near-identical replication of the effect (Figure 7), as well as in the behavioral accuracy from the prior study. The current study extended the physiological findings by also reporting behavioral measures of reaction times for the conditions of context retrieval and contrasted that with recollection-related responses, for both their item and source judgment responses (Figure 3). This revealed reliable differences in how subjects were responding in these instances: participants responded faster to the high confidence hits in item recognition but slower for their more-accurate source judgments, whereas people responded slower at first to the low-confidence hits’ item recognition and faster for their ensuing source judgments.

This extends the ERP findings of Addante et al. (2012) by demonstrating that a) they are not epiphenomenal, b) showing their normative variance as a homogenous effect (Rousselet et al., 2016), and c) indicating that they are behaviorally meaningful in reflecting distinct cognitive processes: retrieving memories of context that are independent of those with recollection. Furthermore, the results revealed that these are independently-replicable findings, as we did so in a different laboratory (CSUSB vs UC Davis), with double the sample size (N=54 vs the original N=25), using different EEG recording hardware (actiCHamp vs BioSemi commercial devices). This replication occurred in a more diverse subject population (CSUSB data was from a federally designated Hispanic Serving Institution comprising 54% of participants as Hispanic, whereas UC Davis was not at the time of the data collection in 2009 and only 24% Hispanic [it has since been recognized as such in 2019]), indicating the simple point that the neurophysiological effects of memory are invariant to such demographic factors not usually studied in college populations.

The behavioral finding of memory performance differences in both speed and accuracy for high- and low-confident hits with correct source memory retrievals indicated that each of the mnemonic conditions for retrieving contextual information of source details was distinct from each other and suggested that they represent at least partially differential neurocognitive processes underlying them, respectively. The full functional significance of the late fronto-central negativity (LCN) remains to be fully determined through future studies of empirical manipulation of systematic variables unavailable in the current investigation, but it continues to be linked thus far to the retrieval of context that lacks the respective processes of both recollection and item-familiarity. This significance is elaborated on below with a prospective account as well as limitations.

Results such as the data reported here and previously (Addante et al., 2012) thus represent important steps in ruling out candidate processes such as recollection and item-familiarity, since the neural correlates were devoid of any indication of the LPC and FN400, respectively, which were instead evident in our several control analyses and those of two prior data sets (R. J. Addante, Ranganath, & Yonelinas, 2012; R. J. Addante et al., 2011). Towards that end, exploratory analyses conducted in the current study identified several related fronto-central-parietal negative-going ERP effects of memory occurring from 800-1200ms for conditions involving retrieval of context (Figure 8), and suggest that the effects are not exclusively limited to those of just low-confidence recognition hits, as related findings of late negative-going ERP effects related to source memory retrieval or context have been reported independently as well (James, Strunk, Arndt, & Duarte, 2016; P. Leynes & Mok, 2020; P. A. Leynes & Mok, 2017; P. A. Leynes & Nagovsky, 2015).

#### 4.1.2 A proposed account of contextual memory retrieval

Putting our observed effects within the context of prior findings, a potential framework can be envisioned to inform how this curious memory situation may arise (i.e. retrieving accurate source memory for items recognized as low-confident hits yet without recollection and above-chance-levels of performance, Figure 2). We propose a working model by which, as a retrieval item is shown on a screen by itself, it doesn’t at first seem familiar (i.e. no FN400, Figure 7). So, one starts to mentally look back in time, engaging a process for searching episodic memory for any context in which to place the item on the screen. During that time (600-800 ms), it is apparent that one cannot recollect the information or the prior event (i.e.: No LPC, Figure 7), which is consistent with the slower reaction times observed for these item judgments (Figure 3). There may still be a general sense of fluency about it from the past, but this would be insufficient on its own to drive accurate performance discriminability, because fluency can often lead to false alarms (Bridger, Bader, & Mecklinger, 2014; Jacoby & Whitehouse, 1989; P. A. Leynes, Landau, Walker, & Addante, 2005; Bruce W. A. Whittlesea, 1993; Bruce W. A. Whittlesea & Leboe, 2003; B. W. A. Whittlesea & Williams, 2001). As the mnemonic search process continues (emerging into the fronto-central negativity, 800-1200) something about the stimuli remains diagnostic for prior exposure, presumably that sense of past fluency as one retrieves an aspect of the item’s prior context (i.e.: the study task it was encountered in), though the information on its own is not enough diagnostically to provide a definitively high confidence response (i.e.: low confidence source-correct).

This contextual information retrieved about the task is enough to be familiar- which is also consistent with the faster reaction times we observed for this condition’s source judgment (Figure 3). In making memory judgments, one can then draw upon that sense that the context seems familiar (i.e.: context familiarity, or contextual fluency), in order to derive the logical inference that the item is likely old, because of the fact that even though you can’t remember it specifically, you do seem familiar with its context, and this basis is sufficient to support an accurate hit in item recognition, albeit with low confidence (item 4 judgments). The item is thus accurately recognized due to the context seeming familiar, or fluent. Though each judgment of item and source are accurately recognized it is via only low confidence, and each of these pieces of episodic information never become bound together into the cohesive representation that we experience both empirically and anecdotally as recollection (i.e. Binding-in-Context model, Eichenbaum et al., 2007; Diana et al. 2008) – which was observed here to exhibit the opposite patterns in both response times and physiological correlates.

Context familiarity would be seen here as a memory process distinct from both those of recollection and item-familiarity processing, consistent with following the framework of the binding-in-context models of memory. This framework provided here for understanding the current results about memory operations both echoes and builds upon the accounts provided in prior work investigating context in memory (R. J. Addante, Ranganath, & Yonelinas, 2012; Montaldi & Mayes, 2010; Reagh & Ranganath, 2018; Yonelinas et al., 2019a), in making the important distinction between item and context processing – particularly distinguishing each for their independence from the episodic memory process of recollection. This account remains to be tested for failures and boundary conditions but is proposed as a working model from which to account for the observed results.

### 4.2 Measures of variance

Recent efforts in the field have noted that measures of variance in both neural time series data and bar graph representations thereof have historically not been very well reported in most cognitive neuroscience discoveries, and many scholars have called for improvements in data reporting (Allen, Erhardt, & Calhoun, 2012; Luck et al., 2020; Rousselet et al., 2016; Rousselet & Pernet, 2011; Weissgerber et al., 2016; Weissgerber et al., 2015). For example, as noted by Rousselet et al., (2016), ‘*using bar graphs [or line plots of neural data] turns a potentially rich patter of results into a simplistic binary outcome,… in which individual differences are ignored*’, and that “*Once bar graphs are replaced by scatterplots (or boxplots etc.), the story can get much more interesting, subtle, convincing or the opposite*”. The current work sought to comply with such recommendations so as to assess the extent of these concerns in the extensively-used ERP measures of recognition memory that had not yet been broadly characterized in such ways (for similar efforts in ERP effects of implicit memory and metacognition, respectively, see (Addante, 2015; Muller et al., 2020).

For the current data of the FN400, LPC, and LFCN, the increased measures of variance and scatterplots of the full raw data indicated that the traditional ERP effects were not contaminated or conflated by the factors of concerns noted above. In all of the effects observed for both behavior and physiology, results exhibited relatively normal distributions and measures of central tendency, while also providing no indications of effects being driven by obvious outliers nor any of the potential concerns raised by Rousselet et al (2016). For instance, what would have been concerning to see in the scatterplots of effects’ variability would have been, for instance, observing two different sub-groups of a bi-modal or tri-modal distribution – but this was not observed and thus provided quality assurance of the validity of the ERP effects of memory. While the ERP effects were revealed to have a range of positive and negative difference values for memory effects, this inherent element of individual ERP data variability has been noted previously in general by Luck (2014) for studies of attention, though has not previously shown specifically for the prominent ERP effects of episodic memory (i.e. FN400 and LPC), and we do so here in a relatively large sized sample of N = 54. One exception to the broad variability seen here of individual subject differences in positive and negative ERP difference scores is a recent ERP study of implicit memory by Addante (2015), which reported all-positive ERP differences for implicit memory measures and which might indicate a more reliable metric for studying individual case studies in small samples such as clinical cases in the future.

There is a broad benefit to the field knowing such basic information about the nature of ERP effects’ variability (as well as behavioral measures’ variability, too), since for instance, these ERP effects of memory have been used internationally to study clinical amnesia patients (Addante, 2015; R. J. Addante, Ranganath, Olichney, et al., 2012; Duzel, Vargha-Khadem, Heinze, & Mishkin, 2001; Duzel, Yonelinas, Mangun, Heinze, & Tulving, 1997; Lehmann, Morand, James, & Schnider, 2007; Mecklinger, von Cramon, & Matthes-von Cramon, 1998; Olichney et al., 2006; Olichney et al., 2002; Olichney et al., 2008; Olichney et al., 2010; Olichney et al., 2000), and thus both the science and society benefit when drawing proper conclusions about data, respectively. The current results represent a step in that direction and provides encouragement for future researchers to adopt similar approaches to data quality measures. Indeed, important future directions for the field is to develop analytic measures for single-subject analyses accounting for individual variability, such as those being developed for oscillatory analyses (Watrous & Buchanan, 2019). Overall, results from individual measures of variability reported here contributes to the field’s further understanding of the traditional ERP effects of memory by providing insight from measures of variance that are not usually reported in prior ERP studies of memory, and though simple in such findings, provides the first comprehensive approach to characterizing the nature of the widely-studied FN00 and LPC effects across measures of standard old-new effects, item recognition confidence contrasts, source memory comparisons, and item+source memory combinations.

### 4.3 Alternatives and Limitations

The full functional significance of the observed effects for the late fronto-central positivity remains relatively novel and hence uncertain beyond the account provided above (section 4.1.2). There are several different accounts which could potentially be used to interpret them for future explorations. One possibility is that it could perhaps reflect a variant of item familiarity-based processing, such as the distinction between absolute (pre-experimental) and relative familiarity (in-the-experiment) of the stimuli (Bridger et al., 2014; Mecklinger & Bader, 2020; A. Mecklinger & R. Bader, 2020). Accurate source memory has been shown to be achieved through reliance on cognitive processes such as item familiarity (P. A. Leynes, Askin, & Landau, 2016; Parks, Murray, Elfman, & Yonelinas, 2011) and unitization (Bader, Mecklinger, Hoppstadter, & Meyer, 2010; Bastin et al., 2013; Diana, Van den Boom, Yonelinas, & Ranganath, 2011; Diana, Yonelinas, & Ranganath, 2008; Parks & Yonelinas, 2015). Indeed, the Source Monitoring Framework (Johnson, Hashtroudi, & Lindsay, 1993; Mitchell & Johnson, 2009) proposed that familiarity can support source judgments in some contexts. However, weighing against such possibilities is that ERP findings for item familiarity, unitization, and item-familiarity contributing to source judgments are found as positive-going ERP difference effects (P. A. Leynes et al., 2016; Mecklinger & Bader, 2020), instead of the opposite pattern of significant negative-going ERP difference effects observed in the current data (Figure 7).

It is also possible that the current results might reflect a form of guessing or implicit memory. In this view, effects may be reflecting conceptual implicit memory contributions to source memory, as has been seen in the form of guessing observed in prior studies of item recognition (Voss & Paller, 2010a, 2010b). However, the fact of their existence corresponding to the high-confidence explicit declarative responses of subjects lends weight against this possibility, as does the above-chance performance of the responses. As recent research is revealing a role for implicit processing fluency making contributions to declarative memory tasks such as cued recall (Ozubko, Sirianni, Ahmad, MacLeod, & Addante, 2020), though, such an account should not be dismissed in entirety as a candidate for the current findings. Indeed, it may be found in future research to be harmonious with the current account provided earlier in discussion, in that the contextual fluency/familiarity may be contributed to by implicit processes giving rise to explicit memory judgments as has been found in other studies of item familiarity and cued recall (Ozubko et al., 2020; Voss, Lucas, & Paller, 2012). As noted earlier in the proposed account of the findings, this familiarity of the contextual may represent various forms of fluency in processing the mnemonic information (Bruett & Leynes, 2015; A. Leynes & Zish, 2012; P. Leynes & Mok, 2020; P. A. Leynes & Addante, 2016; P. A. Leynes et al., 2016; P. A. Leynes, Batterman, & Abrimian, 2019), which has linked negative-going ERP effects for old items to repetition and perceptual fluency. In such an account, it is possible that the negative fluency effects could be times when people are either partially implicitly or explicitly aware of the fluency, and be normative for all old items but in the high-confidence conditions is masked instead by the overwhelmingly positive ERP effects of conscious recollection occurring in the other conditions over-riding the baseline negative effects.

An third possible interpretation of the negative ERP effects for source memory is that they could represent the Late Parietal Negativity (LPN), which is a heterogeneous effect sometimes seen in studies of source memory and other tasks (Johansson & Mecklinger, 2003; Mecklinger, Rosburg, & Johansson, 2016) and which shares similar characteristics to those observed here. The LPN has been associated with a controlled search process for contextual information of episodic memory (Johansson & Mecklinger, 2003; Mecklinger et al., 2016), and is thought to reflect the mental processes of re-constructing the prior episode. The results here are similar to the late parietal negativity (LPN), in that they are negative-going ERP differences that occur late in the epoch and extend to some parietal regions (though observed here and previously to emanate from foci in more fronto-central regions). In their two authoritative Reviews, the LPN was proposed by Mecklinger and colleagues to reflect a reconstructive or evaluative process in memory search when memory features are not fully recoverable, which could indeed be happening in the current study either concurrently with or instead of the model proposed for context familiarity/fluency. Prior work studying the LPN has identified early and late variants of the LPN, with the earlier variant occurring approximately 600 to 1200 ms post stimulus onset and being interpreted as a mnemonic search for context (Herron, 2007), re-activating context-specifying information from an encoded episode (James et al., 2016; Mecklinger, Johansson, Parra, & Hanslmayr, 2007), or with the evaluation of fluency (Herron, 2007; Kurilla & Gonsalves, 2012; Wolk et al., 2004) during later epochs (1200-1900 ms).

The findings by James et al. (2016) of a Cz-centric ERP negativity at approximately 1000 milliseconds that was occurring for source memory retrieval were interpreted as an LPN in that study, but their characteristics could also be interpreted as consistent with the current findings proposed to reflect the familiarity of context. Such interpretations of an LPN could accordingly also potentially be fit to the current results; although, as discussed in the Review by Mecklinger et al (2016) and in Addante et al. (2012), there are several facets which appear to differentiate the current findings from those of the LPN. For instance, the LPN for correct- and incorrect-source has been reported to either be invariant to source accuracy (Herron, 2007) or even to be larger for incorrect source judgments (Wilding, 1999), whereas the current conditions’ effects have been found to be specific for correct source memory (R. J. Addante, Ranganath, & Yonelinas, 2012). The LPN is generally found to be in posterior parietal areas, whereas the current effect was observed in superior central regions of the scalp and distributed to both frontal and central-parietal areas; this topography creates a challenge for the LPN interpretation. Since each effect, respectively, of the ‘LCN’ found here and the LPN reported elsewhere have much room for exploration, the extent to which they may capture partially different, overlapping, or the same constellation of cognitive processes remains to be deciphered in future work.

Since the current experiment was designed only to assess the independent replication of the effect and add behavioral measures of reaction time, the study is unable to fully adjudicate between the range of these possibilities for the functional significance of the negative-going memory ERP effects. Future research specifically designed to empirically manipulate factors on the functional significance of these effects will be needed to make such determinations. What the results can provide, though, is that together, these neural and behavioral findings converge across several studies to reveal a consistent negative ERP effect in fronto-central-parietal sites occurring from 800 to 1200 msec, which exhibited no physiological signs of either item-familiarity (FN400) or recollection (LPC) and yet which exhibited significant above-chance declarative memory performance on source recognition, and did so in a crossover pattern of response speeds when compared to high confidence hits.

### 4.4 Implications for understanding memory organization and processes

The current findings of negative-going ERP effects for successful retrieval of source memory for items that were successfully recognized with low item confidence has now been seen repeated across several studies. First, it was observed in Experiment 1 of Addante et al., (2012), followed by an second internal replication of the effect reported as their Experiment 2 and stemming from the data in a separate study by Addante et al., (2011), and then followed by a third observation of the same ERP pattern in neuropsychological patients lacking hippocampal-based recollection (R. J. Addante, Ranganath, Olichney, et al., 2012). The current results represent a fourth finding of a direct replication of the phenomena, and converges with findings of similar late negative ERP results reported independently for source memory (James et al., 2016). This replication represents external generalizability of the effects across various sample sizes, different demographics, independent laboratories & equipment, and across varying neuropsychological conditions. It thus appears that instances in which people retrieve context of source memory does not necessarily represent de facto evidence of recollection-based processing of that memory.

That implication is important because of the historical role that researchers have given source memory as being a de facto index of recollection when measuring brain activity (Allan & Rugg, 1998; Frithsen & Miller, 2014; Gold et al., 2006; Guo, Duan, Li, & Paller, 2006; Kurilla & Westerman, 2010; Ranganath et al., 2004; M. D. Rugg & Curran, 2007; Slotnick, 2010; Wilding, 2000; Wilding, Doyle, & Rugg, 1995; Wilding & Rugg, 1996; Woroch & Gonsalves, 2010) – and suggests from the current results that the automatic credence previously given to source memory retrieval as a proxy for the process of recollection was a misplaced assumption. Memory theorists have noted for quite some time that source memory is not process-pure as an exclusive measure of recollection (Bowers & Schacter, 1990; Hoffman, 1997; P. A. Leynes et al., 2016; Parks, 2007; Yonelinas, 1999; Yonelinas & Parks, 2007), yet there has nevertheless has been a tendency in the field for neural studies of memory to rely upon contrasts of success vs unsuccessful source memory, or retrieval of context, to infer that recollection is being measured. The current findings make clear that this should be cautioned against, as has been similarly cautioned against for assumptions in measuring item-familiarity (Paller, Lucas, & Voss, 2012; Voss et al., 2012).

This interpretation converges with that of a range of other findings coming from behavioral and neuroimaging approaches that have built a critical mass of support for source memory being supported, at times, by processes other than recollection. For instance, the processes identified by others for making contributions to source memory and retrieval of context have included that of familiarity (Ecker, Zimmer, & Groh-Bordin, 2007; Ecker, Zimmer, Groh-Bordin, & Mecklinger, 2007; Kurilla & Westerman, 2010; Mitchell & Johnson, 2009; Mollison & Curran, 2012; Tsivilis, Otten, & Rugg, 2001; Yonelinas, 1999) and unitization (Diana et al., 2011; Diana et al., 2008; Quamme, Frederick, Kroll, Yonelinas, & Dobbins, 2002). The current results support the interpretations from those findings, in so far as it reveals that recollection is not just merely the retrieval of source memory, since we found clear instances of retrieving source memory that was not linked with neural correlates of recollection (nor of traditional item-familiarity, either).

Together with the current and prior work here, we emphasize that prior approaches of just measuring source memory is not an accurate representative proxy of recollection. Rather, the findings here provide the combined behavioral and physiological evidence to converge with a variety of prior findings affirming that the operational definition of recollection represents the retrieval of mnemonic information of items *bound together* with their information of contextual information (such as source). That view of recollection’s operational definition was initially proposed in the Binding-in-Context (BIC) model by Eichenbaum et al. (2007) and Diana et al. (2007) to be accomplished by the hippocampus, whereas the parahippocampal cortex was modeled to support the independent information of context before achieving recollection-based memories (Diana, Yonelinas, & Ranganath, 2012) – here we provide direct physiological evidence in support of that model and subsequent variants of it (Montaldi & Mayes, 2010; Reagh & Ranganath, 2018; Yonelinas, Ranganath, Ekstrom, & Wiltgen, 2019b). As such, researchers should be advised to be cautious about assuming too much from memory measures of just source memory in the absence of item information, and would be aided to include multiple measures of memory into experimental paradigms, in order to provide the best ways for assuring the measurement of recollection, as opposed to merely capturing its independent constitutive processes (such as item and contextual information processing).

## 5.0 Conclusions

Collectively, the overall patterns of findings directly replicated those reported by Addante, et al. (2012a, 2012b) and indicate that high confidence item recognition coupled with accurate source memory exhibited the neural correlates of recollection and item-familiarity, but that instances of low confidence recognition coupled with accurate source memory did not exhibit the neural correlates of recollection nor familiarity and instead exhibited a later negative-going ERP effect separate in time from the familiarity and recollection-based physiological signals. This revealed that recollection was not present in low confident recognition even in times when accurate source information was retrieved, and furthermore that recollection requires not just having accurate source memory retrieved, as had previously been a general convention for measurement in the field (Ranganath et al., 2004; Wilding, 2000; Wilding & Rugg, 1996).

Thus, in cognitive and neuroscience studies of memory, just because one is comparing correct source memory conditions (e.g. to correct rejections, or to source incorrect conditions), it does not necessarily mean that one is de facto measuring recollection. Just as one can ‘assume too much’ about familiarity-related ERPs for item judgments that can at times represent implicit conceptual fluency (Paller et al., 2012; Voss et al., 2012), or mistake cued recall as representing explicit declarative memory when it can instead be instances of implicit memory (Ozubko et al., 2020), we show here that source memory can be contributed to, and supported by, non-recollective processes that are independent of both recollection and item-familiarity. This conclusion supports the finding from Addante et al (2012) that recollection is independent of source memory and provides physiological support to models that ascribe separable independent roles of context and item processing from recollection-based memory retrieval.

## Acknowledgements

Authors would like to thank Mairy Yousef for helpful assistance with manuscript preparation and literature reviews. Authors thank Drs. Andrew Yonelinas and Andrew Leynes for helpful comments on manuscript drafts.

**This research was supported by the following grants and funding awards**: National Institute of Health Grant 1 L30 NS112849-01 to RJA from National Institute of Neurological Disorders & Stroke; The Mini-Grant Program from the CSUSB Office of Sponsored Research to RJA; the Faculty Assigned Time Grants from the CSUSB Office of Student Research (OSR) to RJA; the Faculty Summer Research Fellowships from the CSUSB Office of Academic Research & the Deans Summer Research Fellowship from the College of Social & Behavioral Sciences to RJA; CSUSB Assigned Time for Exceptional Service to Students Award to RJA from Faculty Senate & Provost; the Faculty-Student Research Grants to RJA, LAS, & AM; the CSUSB Student Success Initiative Innovative Scholars Fund to LAS and AM; and the CSUSB Student Success Initiative Culminating Project Awards to LAS and AM, the Outstanding Graduate Student Award from the CSUSB Dean of Social & Behavioral Sciences to AM, the Outstanding Graduate Student Researcher Award to AM from the CSUSB OSR, and the Outstanding Master’s Thesis Award to AM from the CSUSB Office of Graduate Studies, and start-up research funds & course release time to RJA from Florida Institute of Technology.

## Author Contributions

RJA designed the study, supervised all parts, analyzed data, and wrote the manuscript. AM collected the data and analyzed data. LAS programmed the study.

## Data Accessibility Statement

Data is accessible upon request without restriction, as is any code used to analyze the data-which for the current study involved GUI-based operations in ERPLab. Authors encourage communications from those interested in innovative additional analyses.

## Footnotes

1 Note that for these analyses, source memory confidence was also held constant, utilizing the low-confidence accurate source judgment of ‘4’ in the current paradigm, consistent with the same approach used previously in Addante et al., 2012, and same results were also obtained with collapsing low and high source confidence judgments together.

2 Inspection of the variability of the individual subject data plots identified several subjects (N =6 ) whom performed at- or below-chance levels of source discriminability for accurate source judgments of the item4 responses (Figure 2), which would not have been revealed in a traditional bar graph representation of central tendency (Rousselet et al., 2016). This observation guided us to investigate whether the late fronto-central negativity (LFCN) effects in the ERPs would persist after removal of those subjects from the analysis, which would suggest that the negative ERP effects could be driven in part by spurious factors such as guessing or chance performance. We identified six subjects meeting the criteria for removal in this condition, and results remained unchanged, in that the fronto-central/parietal negative ERP effects persisted in the group of N = 21, indicating that these ERP effects were not being driven by guesses or spurious chance in source memory discriminability, and providing further evidence for the validity of these findings.

